# A theory for how the antigen presentation profile influences the timing of T-cell detection

**DOI:** 10.1101/480301

**Authors:** Alberto Carignano, Neil Dalchau

## Abstract

T-cells are activated when their receptor molecules recognize complexes of MHC proteins bound to peptides on the surface of neighbouring cells. Each T-cell expresses one variant of many possible receptor molecules, which are generated through a partially random process that culminates in approximately 10^7^ possible T-cell receptors. As the peptide sequence bound to an MHC molecule is also highly variable, the optimal strategy of an antigen-presenting cell for displaying peptide-MHC complexes is not obvious. A natural compromise arises between aggressive peptide filtering, displaying a few peptides with high stability MHC binding in high abundance and regularity, and promiscuous peptide binding, which can result in more diverse peptides being presented, but in lower abundance. To study this compromise, we have combined a model of MHC class I peptide filtering with a simple probabilistic description of the interactions between antigen presenting cells (APCs) and cytotoxic Tcells (CTLs). By asking how long it takes, on average, for an APC to encounter a circulating CTL that recognises one of the peptides being presented by its MHC molecules, we found that there often exists an optimal degree of peptide filtering, which minimises this expected time until first encounter. The optimal degree of filtering is often in the range of values that the chaperone molecule tapasin confers on peptide selection, but varies between MHC class I molecules that have different peptide binding properties. Our model-based analysis therefore helps to understand how variations in the antigen presentation profile might be exploited for vaccine design or immunotherapies.

## Introduction

An adaptive immune system is somatically generated during development based on the ability of circulating thymocytes to recognise the protein contents of infected cells. All cells in vertebrates express class I molecules of the Major histocompatibility complex (MHC), comprising the mostly sequence-invariant beta-2-microglobulin (*β*_2_m) and a highly variable heavy chain (HC). In the endoplasmic reticulum (ER), this MHC class I complex binds to protein fragments arising from proteasomal degradation, which are transported from the cytoplasm into the ER by Transporter associated with antigen processing (TAP) molecules. While the specificity of proteasomes and TAP transporters are moderately sequence-dependent, the sheer diversity of the protein sequences of self and viral proteins naturally translates into a large diversity of peptide sequences being transported into the ER. As such, for MHC class I molecules to effectively communicate their internal contents for immune system surveillance, their peptide-binding specificity must be at least moderately promiscuous. It has now recently been shown that the MHC alleles vary in their promiscuity, and that this has a measurable impact on immune response (1).

Cytotoxic T lymphocytes (CTL) detect and destroy infections by recognising antigenic peptides presented on MHC class I molecules at the surface of antigen presenting cells (APC). Each CTL is able to recognise pMHC complexes via their T-cell receptor (TCR), though possess only one TCR type, and so the specificity of the CTL is encoded by its TCR sequence. The TCR is generated in a process of random arrangement of gene segments, known as V(D)J recombination, which creates an enormous diversity of TCRs across T-cell populations. Ideally, there would be sufficiently many T-cells to cover all non-self antigenic determinants, however this would require 10^7^ TCRs to cover 10^13^ antigenic determinants, which naturally requires TCRs to be highly crossreactive, recognising multiple peptides (2).

The formation of the immunological synapse enables TCR to be engaged by peptide-MHC complexes, which can enable CTL to be activated if there are sufficient peptides that the TCR recognise. APCs can present multiple copies of the same peptide sequence, and we term the overall composition of peptides presented as the antigen presentation profile (APP). Given the high degree of crossreactivity between TCRs and peptide-MHC complexes, a compromise might arise in terms of how an APC might optimally select peptides for presentation. For instance, an APP with a small number of distinct peptide sequences in high abundance would maximise the chance that corresponding TCRs will be activated during an interaction. However, an APP with many distinct peptide sequences in lower abundance would increase the number of potential TCR matches, but at the cost of lowering the probability of any one of them leading to activation. Highly relevant to this question is the chaperone molecule tapasin, which skews the APP towards containing more high affinity peptides overall (3–6). Loss of tapasin leads to poor peptide selection, resulting in overall lower cell surface abundance of pMHC. Consequently, tapasin-deficient organisms have altered responses to viral clearance (7–9). It is therefore not surprising that some viruses and tumours down-regulate tapasin to evade immune detection (10). Also of relevance to presentation strategy is the intrinsic binding properties of the MHC haplotype, which have been shown to vary considerably (1).

Theoretical work has already been instrumental in understanding how T-cell responses depend on the biochemical parameters of specific TCR-pMHC interactions. It is now well established that an intermediate rate of TCR-pMHC unbinding (*k*_off_) can provide a sufficient duration for down-stream signalling without preventing additional receptor interactions (11, 12). Computational models of broader scope have helped to interpret how CTL killing depends on cell density within lymph nodes (13, 14). Stochastic models have helped to formulate a probabilistic treatment of T-cell activation in terms of the APP: for instance, (15) modelled the idea that infections are detected when the sum of the stimuli received by a T-cell from the interactions between its TCR and the pMHCs on the APC, exceeds a given threshold. The ability of T-cells of distinguishing between self and non-self is a consequence of a higher copy number of foreign peptides in comparison with self on the APC surface (*probabilistic recognition*). In particular, increasing the copy number of foreign peptides on the APC does not increase the average number of stimuli per T-cell, but the variance, suggesting that exceeding the activation threshold is possible only for high value of foreign peptide copy number (16). However, these theories have not yet been applied to specific MHC haplotypes, and do not consider realistic APC surface presentations. Therefore, here we sought to apply probabilistic models to the analysis of APPs that arise in different MHC haplotypes, and determine how they affect the timing of the CTL response. In order to make this relevant to known MHC molecules, we have made use of the extensive literature on peptide-MHC binding prediction, but also a kinetic model of peptide-MHC loading and presentation (5) to generate plausible antigen presentation profiles.

## Results

### Predicting the cell surface abundance of multiple peptides

Measuring the APP is made challenging by the sheer diversity of peptides, their low copy numbers, a lack of sufficiently many peptide-specific probes, cell-cell variability, and possibly many other factors. Therefore, we base our analysis on the T-cell activation capability of different APPs on computational models, but focus on well characterised MHC class I molecules. As the rate of peptide-MHC dissociation is the major determining factor of cell surface abundance, we sought to create a model that simulates combinations of peptides that differ only in their MHC dissociation rates. This assumes that i) distinct peptide sequences are in equal intracellular abundance, and ii) the association rate of peptide-MHC is homogeneous. To assign dissociation rates to the peptides, we made use of the extensive literature on peptide binding predictors (see Methods).

#### MHC class I alleles differ in their binding repertoires

The BI-MAS algorithm (17) was used to approximate the distribution of dissociation rates of five different HLA alleles (HLA-A*02:01, HLA-B*08:01, HLA-B*27:05, HLA-B*44:03 and HLA-B*58:01). For each allele, we applied the algorithm to a large number of peptides of either 9 or 10 amino acids in length (see Methods for further details). We found that the HLA alleles differed considerably in their distributions (Figure 1). HLA-B*27:05 was found to be highly promiscuous, binding over 4% of peptides with at least moderate stability (Table 1). All other alleles had fewer than 1% of peptides in this category, with HLA-B*08:01 predicted to bind as few as 0.04% with moderate stability.

**Fig. 1.**
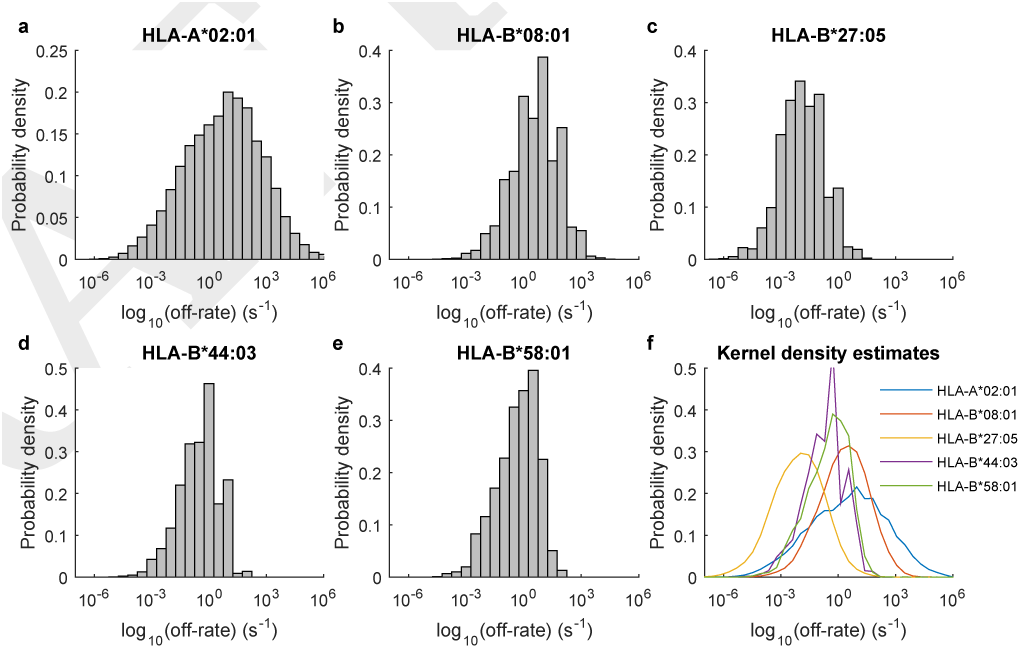
Allelic variation in the distribution of peptide-off-rates from HLA molecules. (**a–e**) The BIMAS algorithm was used to predict the rate of dissociation for complexes of human protein-derived peptides and HLA molecules. Peptides of length 9 and 10 were included in the analysis, and HLA alleles considered are as indicated on each panel. (**f**) For each distribution, a kernel density estimate was obtained using the kde function on MatlabCentral (18), to facilitate comparison of each allele.

**Table 1.** Statistics of peptide off-rate distributions. Reported are the inferred means and standard deviations of the base-10 logarithm of peptide off-rate distributions for HLA molecules. The final column shows the percentage of off-rates that were more stable than 10^−4^ s^−1^

| Allele | $\log_{10}(\text{Mean})$ | Std. | $< 10^{-4} \text{ s}^{-1}$ |
| --- | --- | --- | --- |
| HLA-A*02:01 | 0.73 | 2.00 | 0.86% |
| HLA-B*08:01 | 0.51 | 1.22 | 0.04% |
| HLA-B*27:05 | −1.88 | 1.19 | 4.35% |
| HLA-B*44:03 | −0.55 | 1.05 | 0.25% |
| HLA-B*58:01 | −0.35 | 1.07 | 0.16% |

#### Simulation of the cell surface peptide distribution

Having established considerable variation in the distribution of peptide dissociation rates from different HLA alleles, we sought to characterise how this influences the shape of the distribution of peptides presented at the cell surface, the APP. To provide quantitative predictions of the cell surface abundance of each peptide, the peptide filtering model (5) was simulated for sets of peptides sampled from the distributions in Fig. 1. This allowed us to take into consideration both the peptide dissociation rate from a given MHC class I molecule and also the competition from other peptides for MHC class I loading. Furthermore, it was possible to simulate and therefore predict the consequences of perturbing the MHC class I system, such as in the case of tapasin-deficient cells.

Simulations were carried out for HLA-B*44:02 and HLA-B*27:05 molecules, as these were previously characterised as tapasin-dependent and tapasin-independent alleles in experimental studies (3). In (5), tapasin-dependency was predicted to be attributed to enhanced peptide-MHC association, and in (6) the hypothesis has been refined to rely on a transition from a peptide-receptive to a peptide-nonreceptive conformation (i.e. closing). Since we had used the model in (5), simulations of each allele used different rates of peptide-MHC association (HLA-B*44:02 – 3.1773 × 10^−11^ molecule^−1^ s^−1^; HLA-B*27:05 – 1.9446 × 10^−9^ molecule^−1^ s^−1^). In each case, the off-rates of 10,000 peptides were sampled from lognormal distributions as parameterised in Table 1 (Fig. 2a,d). For HLA-B*44:02, we used off-rates for the closely related HLA-B*44:03 haplotype. Simulations of tapasin-competent and tapasin-deficient cells were carried out for both HLA molecules.

**Fig. 2.**
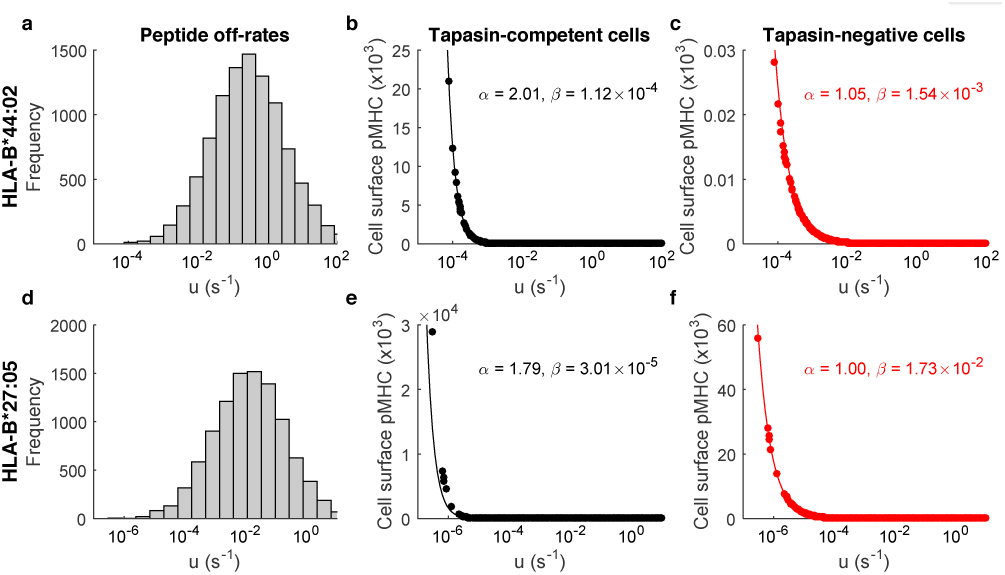
Approximation of the MHC class I peptide filtering model with a power-law function. The peptide filtering model (5) was simulated for 10,000 peptides sampled from log-normal distributions corresponding to (**a-c**) HLA-B*44:02 and (d-f) HLA-B*27:05. (**a**,**d**) Histogram shows the distribution of peptides sampled for each MHC class I allele. (**b**,**e**) Simulations of tapasin-competent cells, for peptides as shown in panels a and d respectively. (**c**,**f**) Simulations of tapasin-negative cells. In panels b, c, e and f, the parameters of the power-law function *βu*^*-α*^ that best fit the simulation of the peptide filtering model are shown.

The effect of *peptide filtering* is clearly visible in Figure 2, as ER distributions of peptides with mode values between 10^−2^ and 10^0^ s^−1^ get mapped to cell surface distributions which are highly skewed towards peptides with low off-rates. The simulations revealed strongly tapasin-dependent presentation for HLA-B*44:02, with two peptides presented at approximately 10^4^ copies each in tapasin-competent cells (Fig. 2b), and *≈* 60 copies each in tapasin-deficient cells (Fig. 2c), a fold change of *≈*2, 000. For HLA-B*27:05, a strongly dominant peptide was presented at 3.7×10^7^ copies in tapasin-competent cells (Fig. 2e) and at *≈*6×10^4^ copies in tapasin-deficient cells (Fig. 2f), a fold change of *≈* 500, indicating tapasin-dependency that is less than that of HLA-B*44:02. It is important to note that the simulated numbers of peptide-MHC complexes at the cell surface are considerably larger than what has been observed experimentally. For instance, it is reported in (19) that the cell surface abundances of specific peptides are in the range 0–100, whereas our simulations indicate abundances as high as *O*(10^7^) (Figure 2). This discrepancy most likely arises because the original model was not calibrated against absolute numbers of molecules, but instead against fluorescence measurements that were assumed proportional to cell surface abundance (5). Accordingly, while the dynamics of the model reproduced experimental observations well, the absolute scale of the model remains uncertain, and simulations should be considered as accurate only up to a multiplicative factor. While the output of the model could be rescaled for its usage in this study, this would not influence the qualitative nature of the results presented. Therefore, for consistency, we leave the peptide filtering model in its original scale.

#### Characterising the cell surface repertoire using a power-law function

While the peptide filtering model (5) provides a biochemically-based simulation of peptide selection, it does not provide a succinct description of the APP as a function of pMHC off-rate. However, we found that the equilibrium cell surface abundance of pMHC could be well approximated by the power-law function

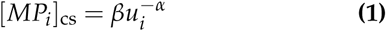

where *u*_*i*_ is the dissociation rate of peptide *i* from the MHC class I molecule (Fig. 2). The major advantage of using this simplified model was that it enabled us to determine the effect of the essential features of the APP on T-cell activation. The parameter *α* quantifies the degree of peptide filtering, which describes the extent to which peptides are selected according to their rate of dissociation from MHC class I. A value of 0 would indicate no dependence on *u*_*i*_, with higher values indicating stronger selection for more stable peptides. Previous theoretical analysis (5) suggested that peptide filtering is at best proportional to 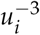 in tapasin-competent cells (*α* = 3), and proportional to 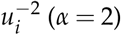 in tapasin-negative cells. The parameter *β* is a scale factor, and corresponds to the average cell surface abundance of a peptide with off-rate 1 s^−1^. It incorporates the supply of peptide into the ER but also incorporates the overall competition between peptides. In this study, we ignore the effect of differential peptide supply as mediated by protein abundance, proteasomal degradation and TAP binding, and so a peptide non-specific value of *β* can be applied to all peptides.

To quantify how the APP differs between alleles and between tapasin-competent vs. tapasin-deficient cells, we inferred values of *α* and *β* in each of the four scenarios of Figure 2 (see Methods for details). The degree of peptide filtering was 2.01 for tapasin-competent cells expressing HLA-B*44:02, though only 1.79 for tapasin-competent cells expressing HLA-B*27:05, emphasising the tapasin-dependency of HLA-B*44:02. In tapasin-negative cells, the degree of filtering was low for both alleles (HLA-B*44:02 – 1.05; HLA-B*27:05 – 1.00). In all cases, the power-law function provided a good approximation of the peptide filtering model for all peptides that had abundance ≥ 1. Crucially, the change in the shape of the distributions was well described by the different values of *α*. In tapasin-competent cells, the large *α* was consistent with high presentation of the most highly stable peptides, at the cost of filtering out lower stability peptides (Fig. 2b,e). In tapasin-negative cells, low *α* was consistent with overall lower presentation, but also a greater number of peptides with moderate presentation.

Since the number of distinct peptide sequences that might be present in the ER at any one time is very large, and beyond what is feasible to simulate with this dynamical model, we performed a number of simulations in which the number of distinct peptides sampled from the off-rate distributions was varied. In doing so, we found that if more than approximately 10,000 peptides were used, the expected deviation from the mean, representative of a larger sample, was below 1% for both *α* and *β*, and for both tapasin-competent and tapasin-negative cells (Fig. S1).

### Probabilistic model of APC-CTL interactions

To interpret how the distribution of peptides presented by MHC molecules on antigen presenting cells influences T-cell recognition, we developed a probabilistic model. The model describes interactions between CTLs and a single APC to determine how differences in the distribution of cell surface peptide-MHC might affect the timing of T-cell activation. For clarity, the parameters of the model are summarised in Table 2, but we introduce many of these quantities as we describe the model. Dendritic cells are thought to interact with as many as 5,000 T cells per hour *in vivo*, which is approximately 1 per second (19). Therefore, the time unit of this model can be conveniently thought of as 1 second. The same study (19) measured the average contact duration to be 3.4 minutes, suggesting that on average there are approximately 200 T-cells in contact with an APC at any one time. We may therefore formulate a simple model for the interactions between a single APC and multiple CTLs as

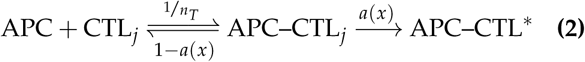

where *n*_*T*_ is the assumed number of CTLs and APC–CTL^*∗*^ represents an activated T-cell bound to the APC (Fig. 3). The activation rate *a*(*x*) is a function that depends on the current surface distribution of the APC. We use the variable *x* to denote a random sample of the cell surface distribution, corresponding to the peptide-MHC molecules that are scanned in an immunological synpase. The activated state is considered to be *absorbing* in this model, which means that it is the final state considered. Moreover, the probability of T-cell activation is always positive here, which assumes that there is always a peptide on the surface for which at least one T-cell can recognise. Related to this, we note that negative selection, the deletion of T-cells that react with self peptide, is not explicitly considered here.

**Table 2.** **Definitions of modelling quantities used throughout.**

| Variable | Description | Value(s) |
| --- | --- | --- |
| $u_i$ | Dissociation rate of a peptide-MHC complex | Variable ($> 10^{-6} \text{ s}^{-1}$) |
| $\beta$ | Relative supply of each peptide | Inferred (HLA/Tpn-dependent) |
| $\alpha$ | Skewness of peptide selection | Inferred (HLA/Tpn-dependent) |
| $[MP_i]_{cs}$ | Number of peptide-MHC complexes with peptide $i$ | Eq. 1 |
| $P_T$ | Number of peptide-MHC complexes on the cell surface | $\sum_i [MP_i]_{cs}$ |
| $T_T$ | Number of TCRs on the surface of a CTL | 1000 |
| $K_D$ | Half saturation constant | 10 |
| $N_P$ | Number of distinct peptide sequences that can be presented on MHC molecules | |
| $n_{cog}$ | Number of distinct peptide sequences that are cognate for a TCR | |
| $n_{scan}$ | Number of pMHC-TCR bonds | Eq. 5 |
| $n_{diff}(x)$ | Number of distinct peptide sequences in a sample ($x$) of pMHC molecules presented on the APC surface | ($\leq n_{scan}$) |

**Fig. 3.**
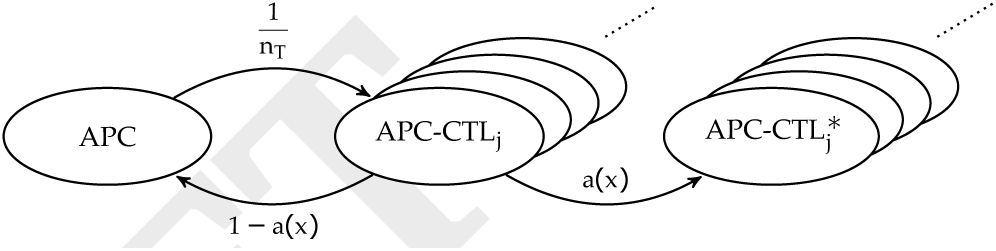
Markov process model of APC-CTL interactions. The binding of an APC to a CTL is considered to take place once per time unit. Therefore, we normalise against the assumed number of CTL, *n*_*T*_. Upon formulation of the immunological synpapse, the CTL is either activated with probability *a*(*x*), which is a function of the APP *x*, or dissociates with probability 1 - *a*(*x*). Multiple APC-CTL complexes are shown to illustrate that there are multiple CTLs that might bind, each occurring with frequency 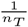.

**Fig. 4.**
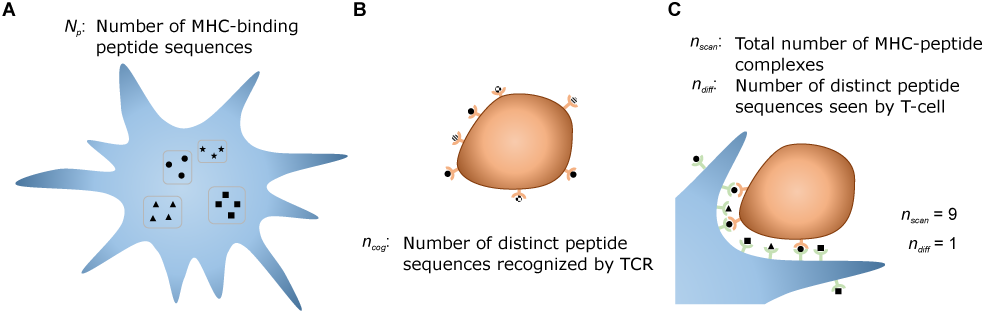
Quantifying pMHC-TCR interactions. Shown are a series of diagrams that illustrate how model parameters are defined. An antigen-presenting cell is shown in blue, and a T-cell in orange.

The model assumes that, on average, an APC encounters T-cells at a frequency of 1 per time unit. As this is the only time-dependent process, the model can be arbitrarily scaled, and as such we leave the model in terms of this time unit. To further simplify this description so as to enable useful calculations to be carried out, we remove the probabilistic continuous-time nature of T-cell encounters and consider the discrete-time version of the model

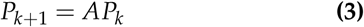

where the state space *P*_*k*_ records the number of copies of each state at time *t* = *k* (*k* = 1, 2, *…*) and the transition matrix *A* describes the transition propensities between each state. We write these as

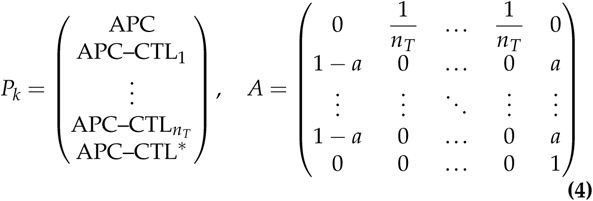

#### Calculating the expected time until first detection

Using our model of APC-CTL encounters, it is possible to derive the expected number of T-cell encounters that are required until a T-cell is activated, which can be used as a proxy for the efficiency of an APC to stimulate a CTL response. We use the fact that the expected number of steps required of a discrete-time Markov chain to reach an absorbing state is given by the sum of the first row of

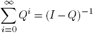

where *Q* is the transient part of the transition matrix, which in this case is given by the first *T*_*n*_ + 1 rows of *A*. As such,

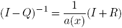

Where

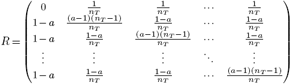

The sum of the top row of (*I* - *Q*)^−1^ is therefore equal to 2*/a*(*x*), which is independent of *n*_*T*_, the number of CTLs. Therefore, in this model, the probability of T-cell activation is determined purely by the probability of activation *a*(*x*).

#### Calculating the probability of T-cell activation

In the model presented here, every time there is a contact between an APC and a CTL, the cell surface distribution is sampled so as to mimic the formation of the immunological synapse and binding of TCRs to surface pMHC complexes. The quantity *a*(*x*) is then used to describe the probability that the CTL is activated from a sample *x* of pMHC complexes at the cell surface. The size of the sample is taken to be a function of the total number of pMHC molecules available for TCR binding (*P*_*T*_ = Σ_*i*_ [*MP*_*i*_]_cs_) and an assumed number of TCRs (*T*_*T*_), as described in (12), as

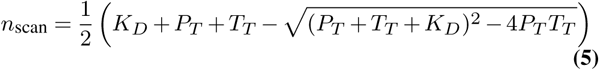

where *K*_*D*_ is the bond dissociation constant, which is the ratio of the off- and on-rates of pMHC-TCR binding. By using this model, we can incorporate the effect of TCR saturation, which will occur if there are very high cell surface levels of peptide-MHC. This imposes a limit on the number of peptide-MHC complexes that are scanned during an APC-CTL contact. As such, simply presenting more peptide-MHC complexes will not lead to more TCRs being engaged.

We use Eq. 5 to define the number of peptides that are sampled by a T-cell that comes into contact with the APC. This means that we have reinterpreted the *K*_*D*_ parameter to be an *averaged* dissociation constant over the different peptide sequences that are bound to the MHC molecule. As such, *K*_*D*_ is the value that equates

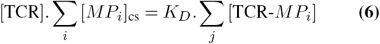

While the value of *K*_*D*_ is likely to be inaccurate for any specific pMHC-TCR interaction, the use of the equation represents the simplest way to describe the potential limitation of TCRs in the immunological synapse, which will become important as total APC presentation of pMHC increases.

To calculate the activation probability from a sample of *n*_scan_ pMHC complexes, we make some important simplifying assumptions. First, we assume that each T-cell may be activated by a fixed number of *cognate* peptides *n*_cog_. Second, in a sample *x* of the APP, *n*_diff_(*x*) different peptides are seen by the T-cell. In this way, the activation probability *a*(*x*) follows a hypergeometric distribution over *N*_*P*_, the total number of peptides that might be presented. Since encountering one cognate peptide is thought to be sufficient to activate a T-cell (20), albeit via multiple TCR engagements (21), we formulate an approximate activation probability on the event that at least one cognate peptide is encountered. As such the probability of activation is 1 the probability of not encountering a cognate peptide, i.e.

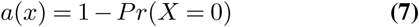

where *X* is a hypergeometric distribution with parameters (*N*_*P*_, *n*_cog_, *n*_diff_(*x*)).

Based on this definition for *a*(*x*), it remains for us to determine *n*_diff_, the number of different pMHC seen by the TCRs. Since several copies of each peptide are present on the surface of the APC, *n*_diff_ will be less than the total number of peptides scanned by the T-cell (*n*_scan_). For example, peptides with high affinity for one of the expressed MHC molecules will likely have higher copy numbers than lower affinity ligands. Moreover, the number of different peptides is huge (*N*_*P*_ > 10^15^) (22). Hence, an exhaustive search of all the possible outcomes *n*_diff_ given the number of scanned peptides by the T-cell is computationally intractable and no analytic formula has been derived to our knowledge. However, one can compute its expected value as a function of the surface multiplicity of each peptide *n*_*i*_ (defined by Eq. 1) and *n*_scan_. To obtain a probabilistic expression for *n*_diff_, we introduce the index variable *I* defined as:

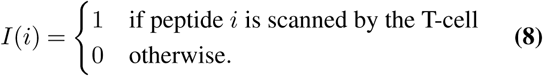

Using the index variable *I*, we can find an expression for the expected number of *n*_diff_ as:

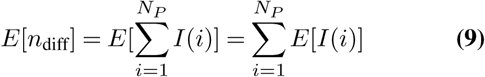

The expected value of *I* (*i*) is by definition equal to 1 – the probability that peptide *i* was not scanned. This is equal to:

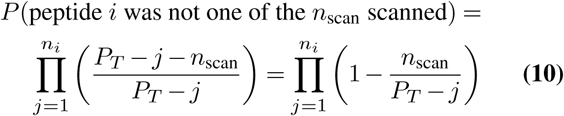

Hence, combining Eq. 9 and Eq. 10, we obtain the expected value of *n*_diff_ as:

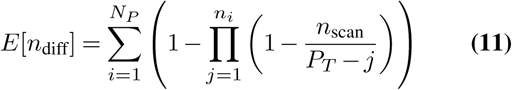

The probability of activation *a*(*x*) can also be obtained by using *E*[*n*_diff_] instead of *n*_diff_ in Eq. 7.

To make the calculation of the expected time until first detection more computationally efficient, we grouped peptides together according to their cell surface multiplicity. By defining *k*_*i*_ to be the number of peptides with cell surface multiplicity *i*, and as such 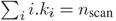, we can rewrite Eq. 11 as

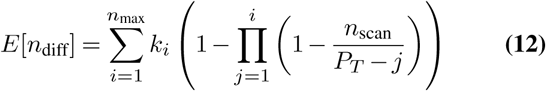

where *n*_max_ is the highest multiplicity observed. To summarise the procedure for calculating the expected time until first detection, we present the steps in Algorithm 1.

##### Algorithm 1 Calculating the expected time until first detection

1: Sample peptides with log_10_(*u*_*i*_) ∼ 𝒩 (*µ, σ*^2^) for *i* = 1, *…, N*_*P*_.

2: Quantify their cell surface abundance bound to MHC class I, using the peptide filtering model (5), or the power-law function (Eq. 1).

3: Round each abundance value to the nearest integer, producing a set {*P*_*i*_∈ ℤ^*∗*^: *i* = 1, *…, N*_*P*_}.

4: Determine the number of pMHC-TCR contacts, *n*_scan_, using Eq. 5.

5: Determine the expected number of different peptides presented using Eq. 11.

6: Evaluate the activation probability *a*(*x*) using Eq. 7 with*n*_diff_ = *E*(*n*_diff_).

7: Evaluate the expected time as 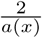.

### The distribution of peptides on an APC modulates the expected time until first productive T-cell encounter

To determine how the timing of T-cell activation depends on the distribution of peptide off-rates for an MHC class I molecule, we used the power-law description of cell surface peptide-MHC abundance within the probabilistic model of T-cell detection (Fig. 3). Samples of peptides were drawn from log-normal distributions with varying mean (*µ*) and standard deviation (*σ*) parameter values (Fig. 5a). For each pair of values, the cell surface abundance was then calculated for a varying degree of filtering (*α*) and peptide supply (*β*), and then used to calculate the expected time until first detection using the Algorithm 1.

We found that the time until first detection routinely showed a compromise in the degree of filtering, with fastest detection reached with an intermediate value of *α* (Fig. 5b). Our interpretation of these results is that higher filtering raises overall cell surface abundance of high affinity peptides, but at the cost of diversity, with medium and low affinity peptides being filtered out. To understand this more clearly, we consider the extremes. Perfect filtering (*α* → *∞*) would produce a very large abundance of only a single peptide, which would lead to only one class of T-cells being activated (assuming there exists a circulating CTL that reacts with this peptide). Whereas an absence of filtering (*α* = 0) would lead to an equivalent number of each peptide being presented, but each at very low abundance. As total abundance drops with lower *α*, we find that only a small number of total peptides are presented at all when *α* = 0, leading also to a low number of activatable T-cell classes. By increasing the filtering to an intermediate level, total surface abundance increases, but maintains some diversity, leading to a larger number of activatable T-cell classes.

**Fig. 5.**
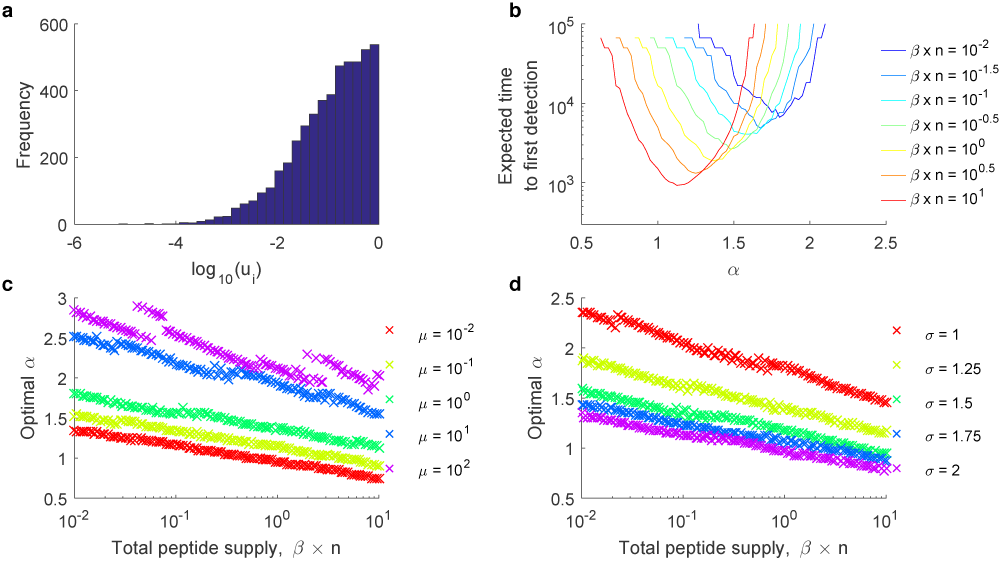
The distribution of peptides on an APC modulates the expected time until first productive T-cell encounter. The expected time until first detection was calculated for peptides sampled from a range of log-normal distributions. (**a**) Shown is an example sample of 10,000 peptide off-rates drawn from a log-normal distribution with mean *µ* = 1 and standard deviation *σ* = 1.5. Note the distribution is truncated to peptides with their off-rates no larger than the mean, for efficiency of the expected time calculation. (**b**) Expected time to first detection as a function of *α* and *β*, specifically for the case in (a). (**c**) Shown is the value of *α* that minimises the time until first detection, as a function of peptide supply (quantified as *β × n*, where *n* is the number of peptides sampled), but also for varying *µ* (*σ* fixed at 1.25). (**d**) As in (c), but this time varying *σ* (*µ* fixed at 1).

As the supply of peptide was decreased, the optimal degree of filtering increased, but the expected time until first detection lengthened (Fig. 5b). Our interpretation is that in low supply scenarios, additional filtering can compensate for a lower total number of cell surface complexes, but will naturally result in fewer distinct peptides, and therefore fewer numbers of activatable T-cell classes.

We found that the optimal degree of filtering also varied with the underlying distribution of peptide off-rates. As the mean of the off-rates became lower (peptides bind MHC more stably), the optimal *α* decreased (Fig. 5c). This is because lower off-rates are presented in higher abundance, and so less filtering is required to maintain a sufficiently high total abundance to nearly saturate the TCRs. Similarly, increasing the standard deviation of the off-rates decreased the optimal *α* (Fig. 5d), as this increases the number of high stability peptides in the sample.

### Tapasin can accelerate or decelerate T-cell detection, depending on HLA allele

To determine how enhanced peptide filtering, conferred by the chaperone molecule tapasin, might influence the timing of T-cell detection, we applied our probabilistic model to two HLA alleles. Differences in the tapasin-dependency of HLA-B*27:05 and HLA-B*44:02 was previously modelled by changing the inrinsic association rate of peptide-MHC binding (parameter *b* in (5)). A low *b* (HLA-B*44:02) translates into strong tapasin-dependency, with few peptide-MHC molecules being detectable at the cell surface in tapasin-negative cells (3, 5). A high *b* (HLA-B*27:05) permits sufficient peptide loading in tapasin-negative cells for detectable cell surface presentation, but in tapasin-competent cells leads to some peptides being loaded on tapasin-unbound molecules, circumventing the potential benefits of tapasin, and thereby reducing tapasin-dependency overall (3, 5).

We carried out simulations for HLA-B*27 and HLA-B*44, using the off-rate distributions quantified in Table 1, varying the on-rate *b* over a large range, and comparing tapasin-competent with tapasin-negative cells. In tapasin-negative cells, increasing the on-rate *b* always sped up T-cell detection (Figure 6). At the inferred value for *b* (from (5)), HLA-B*27 had a surface distribution which produced efficient T-cell detection, such that increases in this rate provided no additional benefit. In contrast, the inferred value of *b* for HLA-B*44 did not lead to any peptide-MHC complexes at the surface of tapasin-negative cells. In tapasin-competent cells, we found that the expected time to first detection strongly differed between the HLA alleles. We found that tapasin expression led to less efficient T-cell detection for HLA-B*27, though more efficient T-cell detection for HLA-B*44 (Figure 6). Since tapasin expression increases peptide filtering (parameter *α*), we can use the analysis of Figure 5b to explain how this arises. HLA-B*27 presumably corresponds to the situation exemplified by the red trace in Figure 5b, with an optimal *α* near 1, and declining detection with increased *α*. On the other hand, HLA-B*44 presumably corresponds more to the situation exemplified by the blue trace, where the optimal *α* is closer to 2, and less peptide filtering leads to very slow T-cell detection. We suggest that the shift in the optimal degree of filtering of these two HLA alleles results from a shift in the means of the off-rate distributions. HLA-B*27 has a log_10_ mean of -1.88 and HLA-B*44 has a log_10_ mean of -0.55 (Table 1), which we would expect to increase the optimal degree of filtering (Figure 5c).

**Fig. 6.**
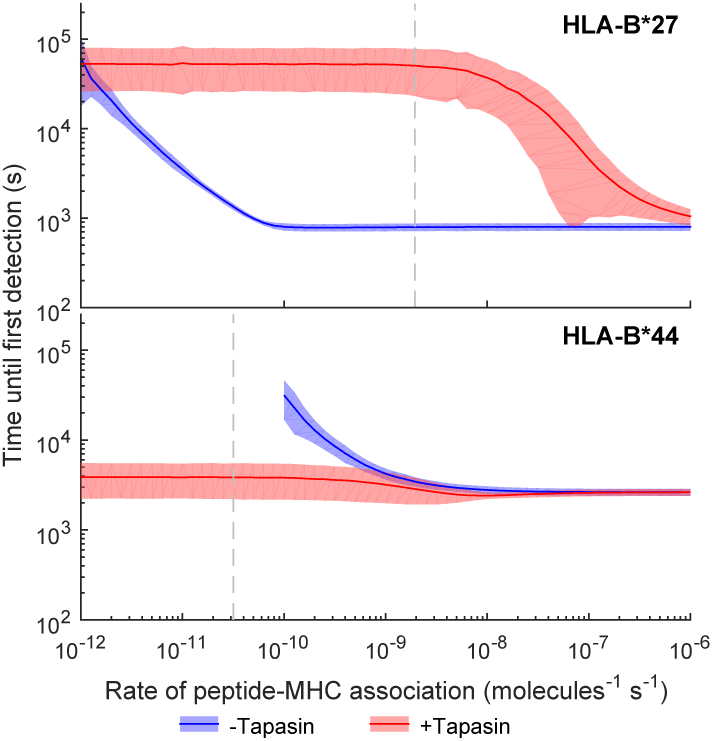
Tapasin can accelerate or decelerate T-cell detection, depending on HLA allele. The rate of peptide-MHC association (*b*) was varied in simulations of HLA-B*27 (top panel) and HLA-B*44 (bottom panel). Compared are simulations of tapasin-competent (red) and tapasin-negative (blue) cells. The expected time until first detection is quantified according to Algorithm 1. Simulated cell surface abundance incorporated the allele-specific peptide-MHC on-rates established in (5), with *b* = 1.9446 × 10^−9^ molecule^−1^ s^−1^ for HLA-B*27 and *b* = 3.1773 × 10^−11^ molecule^−1^ s^−1^ for HLA-B*44. These rates are indicated by the dashed lines.

## Discussion

Due to the combinatorial nature of antigen presentation, it has been difficult to assay for how T-cell activation depends on the genetic variations in MHC class I. By combining current quantitative understanding of intracellular peptide-MHC binding and cell surface presentation with pMHC-TCR binding in the immunological synpase, the theory presented illustrates how changes in the antigen presentation profile can influence the timing of T-cell responses. Accordingly, the theory provides a perspective on how changes in antigen presentation, as encountered during viral infection, might affect an immune response. During viral infection, it is known that interferon-*γ* (IFN*γ*) release stimulates enhanced expression in the MHC locus, leading to enhanced expression of MHC class I/II and other components of the PLC such as tapasin and TAP (23), which all increase overall antigen presentation (24). In contrast, many viruses interfere with the antigen presentation pathway, reducing or at least altering the composition of peptide-MHC molecules presented at the cell surface (25). The HIV protein Nef is known to target pMHC complexes in post-ER compartments (26), while human cytomegalovirus impairs formation of the PLC and reduces tapasin expression (10).

We found that changes in the antigen presentation profile (APP) of antigen presenting cells can lead to a non-intuitive impact on the expected time until first productive T-cell encounter. Depending on the supply of peptide-MHC molecules intracellularly (parameter *β*), efficient T-cell detection could be achieved optimally by either aggressive or weak peptide filtering, depending on the MHC alleles. When peptide supply is low, more aggressive filtering is needed to overcome low numbers of peptide-MHC molecules arriving at the cell surface. Whereas, when peptide supply is higher, the cell surface becomes saturated. In this case, weaker peptide filtering results in greater diversity in the APP, and therefore leads to a greater number of T-cells having the potential to be activated by the APC. This observation could help to explain why MHC class I alleles have evolved with variable dependency on the chaperone protein tapasin (3), which enhances peptide filtering (4–6).

A natural extension of our results could be obtained by incorporating some of the ideas developed in (16), which considers the more general scenario of *probabilistic recognition* of infections. Whereas our analysis computes the probability that a T-cell encounters at least one cognate peptide, the framework introduced by (16) considers partial activation by non-cognate peptides. Our model can be compared with their results, for T-cell clonotype *j*, by assigning *W*_*j*_ = 1 and *W*_*i*_ = 0 for *i* ≠ *j*, and setting an activation threshold of 1. Our analysis could be extended to account for higher activation thresholds, since Eq. 7 follows a hypergeometric distribution, though this would not change our conclusions. In fact, our analysis corresponds to the case where T-cell *mistakes* are negligible. And according to (16), this could always be made true for large populations of cognate peptides on the APP (*z*_*f*_ in their study) by adjusting the threshold of activation accordingly.

Similarly to (16), we did not consider the mechanism of negative selection explicitly in our study. However, this effect could be easily added to our framework by adjusting the transition matrix *A* in Eq. 4. Rows corresponding to T-cells that cannot be activated by an APC only presenting self-peptides are set to [100···0]. The expected time for detection then becomes 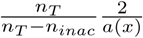, where *n*_*inac*_ is the number of T-cells that do not respond to the APC, which amounts to a simple scaling factor. Therefore, consistent with (15), T-cell activation dynamics are not qualitatively affected by negative selection, although the activation threshold is reduced. Furthermore, our analysis into the effect of tapasin remains unaltered, since the quantity *a*(*x*) is independent of negative selection.

By combining established theories of antigen presentation and T-cell stimulation, we have produced predictions of immune system functioning that we hope can be tested experimentally. As we have simulated established HLA alleles directly, those same alleles could be incorporated into experimental analyses. However, murine models offer more potential for obtaining quantitative estimates of APPs in infected or uninfected organisms, and downstream T-cell responses. New methods based on mass spectrometry are beginning to enable high throughput quantitation of cell surface peptide-MHC, sorted by peptide sequence (27), offering the means to characterise the APP in detail, and over time. By measuring the timing of T-cell responses in a corresponding range of MHC backgrounds, it will soon be possible to establish whether the composition of the APP, rather than simply the presence of a handful of immunodominant epitopes, is a more representative determinant of T-cell response times.

## Materials and methods

### Off-rate prediction using BIMAS

The dissociation rates of five HLA class I alleles were characterised by applying the BIMAS algorithm to a large set of peptides. The peptides were generated from human protein sequences obtained from UniProtKB (release 2010_08). All subsequences of length 9 and 10 were extracted from the sequences of the proteins across the 24 chromosomes, generating a total of 22,160,455 peptide sequences. The dissociation rates were computed in Matlab, following the methodology described in (17).

### Inferring the power-law parameters from simulated cell surface pMHC abundance

For a given distribution of peptide off-rates according to log_10_(*u*) *∼* 𝒩 (*µ, σ*^2^), the values of *α* and *β* are inferred from simulations of the peptide filtering model (5). The power-law function is fitted to equilibrium simulated cell surface abundance of peptide-MHC complexes by first log_10_-transforming the simulated values. Given that log_10_(*βu*^*-α*^) = log_10_(*β*) *- α* log_10_(*u*), we can then identify *α* and log_10_(*β*) using linear regression of the log_10_ transformed equilibrium simulations against log_10_(*u*). We used the *polyfit* function in Matlab, applied to peptides with log_10_(*u*) *< µ* - *σ* to calibrate *α* and *β* in this way. This sub-sample was used to ensure high accuracy approximation of the high affinity peptides.

## ACKNOWLEDGEMENTS

The authors wish to thank Marco Ferrarini for critical reading of the manuscript.

## Supplementary Note A: Supplementary Figures

**Fig. S1.**
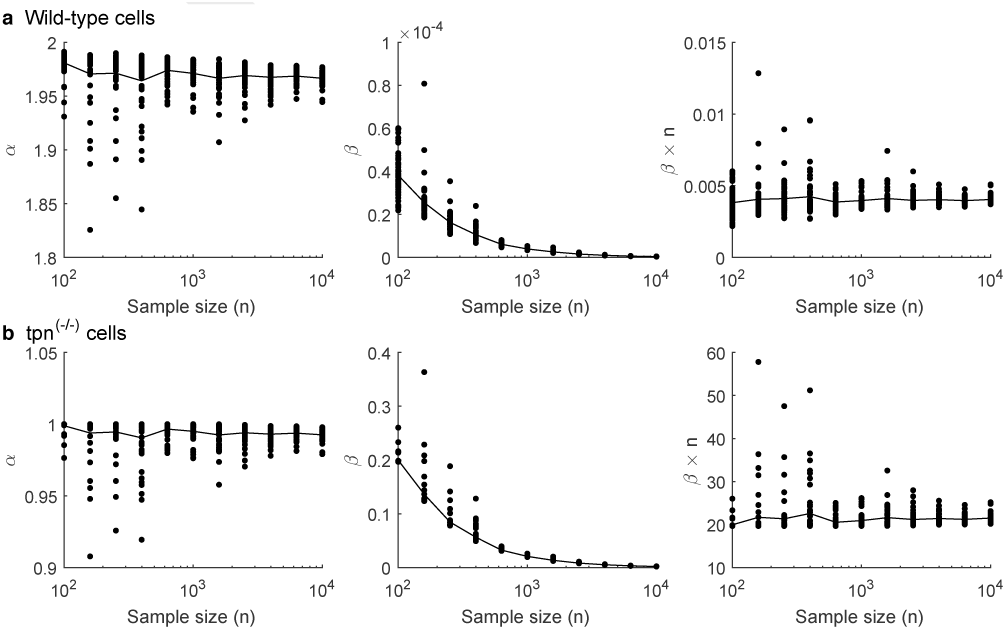
Effect of sample size on relating the MHC class I peptide filtering model to power-law cell surface distributions. The number of distinct peptide sequences (*n*) was varied between 10^2^ and 10^4^, to determine the extent to which under-sampling peptides affects the distribution of peptides presented at the cell surface. This is quantified in terms of the best fit parameter values *α* and *β* of the power-law approximation for cell surface presentation of peptide-MHC complexes. The rightmost panels illustrates that variations in *β* can be compensated for by scaling out the sample size *n*.

